# Universal Panel Testing of Pancreatic Cancer Cases for Cancer Predisposition

**DOI:** 10.1101/195537

**Authors:** Erin L. Young, Bryony A. Thompson, Deborah W. Neklason, Angela K. Snow, Matthew A. Firpo, Theresa Werner, Russell Bell, Nykole R. Sutherland, Ken R. Smith, Justin Berger, Alison Fraser, Amanda Gammon, Cathryn Koptiuch, Wendy K Kohlmann, Leigh Neumayer, David E. Goldgar, Sean J. Mulvihill, Lisa A. Cannon-Albright, Sean V. Tavtigian

## Abstract

**Background and Aims:** Genes associated with hereditary breast and ovarian cancer (HBOC) and colorectal cancer (CRC) susceptibility have been shown to play a role in pancreatic cancer susceptibility. Germline genetic testing of pancreatic cancer cases could be beneficial for at-risk relatives with pathogenic variants in established HBOC and CRC genes, but it is unclear what proportion of pancreatic cancer cases harbor pathogenic variants in these genes.

**Methods:** 66 pancreatic cancer cases, unselected for family history and diagnosed at the Huntsman Cancer Hospital (HCH), were sequenced on a custom 34-gene panel including known HBOC and CRC genes. A second set of 156 unselected HCH pancreatic cancer cases were sequenced on an expanded 59-gene panel (n=95) or with a custom 14-gene clinical panel (n=61). Sequencing data from both sets of pancreatic cancer cases, the pancreatic cancer cases of the Cancer Genome Atlas (TCGA), and an unselected pancreatic cancer screen from the Mayo Clinic were combined in a meta-analysis to estimate the proportion of carriers with pathogenic and variants of uncertain significance.

**Results:** Approximately 8.9% of unselected pancreatic cancer cases at the HCH carried a variant with potential HBOC or CRC screening recommendations. A meta-analysis of unselected pancreatic cancer cases revealed that approximately 10.5% carry a pathogenic variant or HiP-VUS.

**Conclusion:** With the inclusion of both HBOC and CRC susceptibility genes in a panel test, unselected pancreatic cancer cases have a high enough percentage of carriers to rationalize genetic testing for identification of variants that could be further used in cascade testing of healthy relatives to increase HBOC and CRC surveillance measures.

## INTRODUCTION

Over the last few years, massively parallel sequencing converged with targeted capture using array synthesized baits to enable panel testing of most known cancer susceptibility genes.[1–4] These panel tests have since replaced Sanger sequencing of limited sets of syndromic genes, revolutionizing the genetic testing landscape for Hereditary Breast and Ovarian cancer (HBOC) and Colorectal cancer (CRC) predisposition. Before panel testing, the personal and family history of a cancer patient had to be examined carefully to select the most likely syndromic genes. Due to atypical personal or family history combined with the non-discreet nature of cancer susceptibility,[5–10] there were probably many instances where a patient never received the correct test.

Genetic testing of patients with pancreatic cancer is not yet routine, in part because methods for prevention or early detection of pancreatic cancer have limited utility.[11–14] One substantial benefit of genetic testing is the application of preventive measures for carriers of disease susceptibility alleles, notably the at-risk relatives of probands. Pancreatic cancer patients have been reported to carry pathogenic variants in a variety of cancer susceptibility genes, notably HBOC and CRC genes, for which surveillance, medical, and surgical strategies towards prevention or early detection are available. [15–24] Therefore, systematically testing pancreatic cancer cases for pathogenic germline variants in HBOC and CRC genes in order to identify at-risk relatives who would benefit from preventive measures for breast cancer, ovarian cancer, and CRC could be a medically useful application of panel testing. Testing all pancreatic cancer cases for HBOC and CRC susceptibility, however, would only be beneficial if the proportion of carriers among pancreatic cancer cases is high enough to justify that testing.

To estimate the percentage of pancreatic cancer cases that carry variants with potential medical management impact for at-risk relatives, we applied panel testing to independent discovery (n=66) and replication (n=156) sets of pancreatic cancer patients ascertained at the Huntsman Cancer Hospital (HCH) in Salt Lake City, UT, both unselected for family cancer history. To demonstrate generalizability of the results in pancreatic cancer cases, we performed a meta-analysis including published panel tests of unselected pancreatic cancer cases.

## METHODS

### Subjects and Ethics Statement

This study was approved by the Institutional Review Board of the University of Utah. All participants gave written consent, which included DNA sampling for molecular studies and access to medical records.

An initial set of pancreatic cancer cases (n=66) were selected on the minimal requirements of personal history of cancer and having at least two grandparents in the genealogy data represented in the Utah Population Database (UPDB). These patients were screened with a 34-gene custom research panel. Individual family members were then linked to statewide cancer, demographic, and medical information.[25] Ages at diagnosis and family cancer history were obtained from the UPDB after sequencing and variant evaluation. The second set of pancreatic cancer cases (n=169) were selected on the basis of being newly diagnosed pancreatic cancer cases ascertained at the HCH from July 2014 to July 2015, and from December 2015 to April 2017. The pancreatic cancer cases ascertained during the interval July 2014-July 2015 were screened with a 59-gene custom research panel, and the cases ascertained during the interval December 2015-April 2017 were screened with a 14-gene custom clinical panel.

### Next-generation sequencing library preparation and custom targeted capture

For research panel testing, blood-derived genomic DNA (100ng) was sheared using a Covaris S2 instrument (Covaris, Woburn, MA, United States). Genomic libraries were prepared using the Ovation Ultralow Library System (NUGEN # 0329) according to the manufacturer’s instructions. Library enrichment for a 34 or 59-gene custom panel was done with the Roche SeqCap EZ Choice Library (cat# 06266339001) and the SeqCap EZ Reagent Kit Plus v2 (NimbleGen #06-953-247-001) using the manufacturer’s protocol. Individual libraries were combined into pools of 6-12 prior to hybridization, and then super-pooled for up to 96 samples per sequencing lane. Captured libraries were sequenced on an Illumina HiSeq2000 channel using the HiSeq 101 Cycle Paired-End sequencing protocol.

On the strength of preliminary data from this study, HCH began systematically offering clinical panel predisposition testing beginning December 2015, without regard to family history. From December 2015 to April 2017, clinical testing was offered to 73 sequential pancreatic cancer cases. 61 pancreatic cancer cases accepted clinical testing with the 14-gene custom panel was conducted by Invitae. The 12 individuals that declined were in poor health and/or did not see value in undergoing a genetic test. A complete list of genes captured is included in Supplemental Table 1.

**Table 1:**
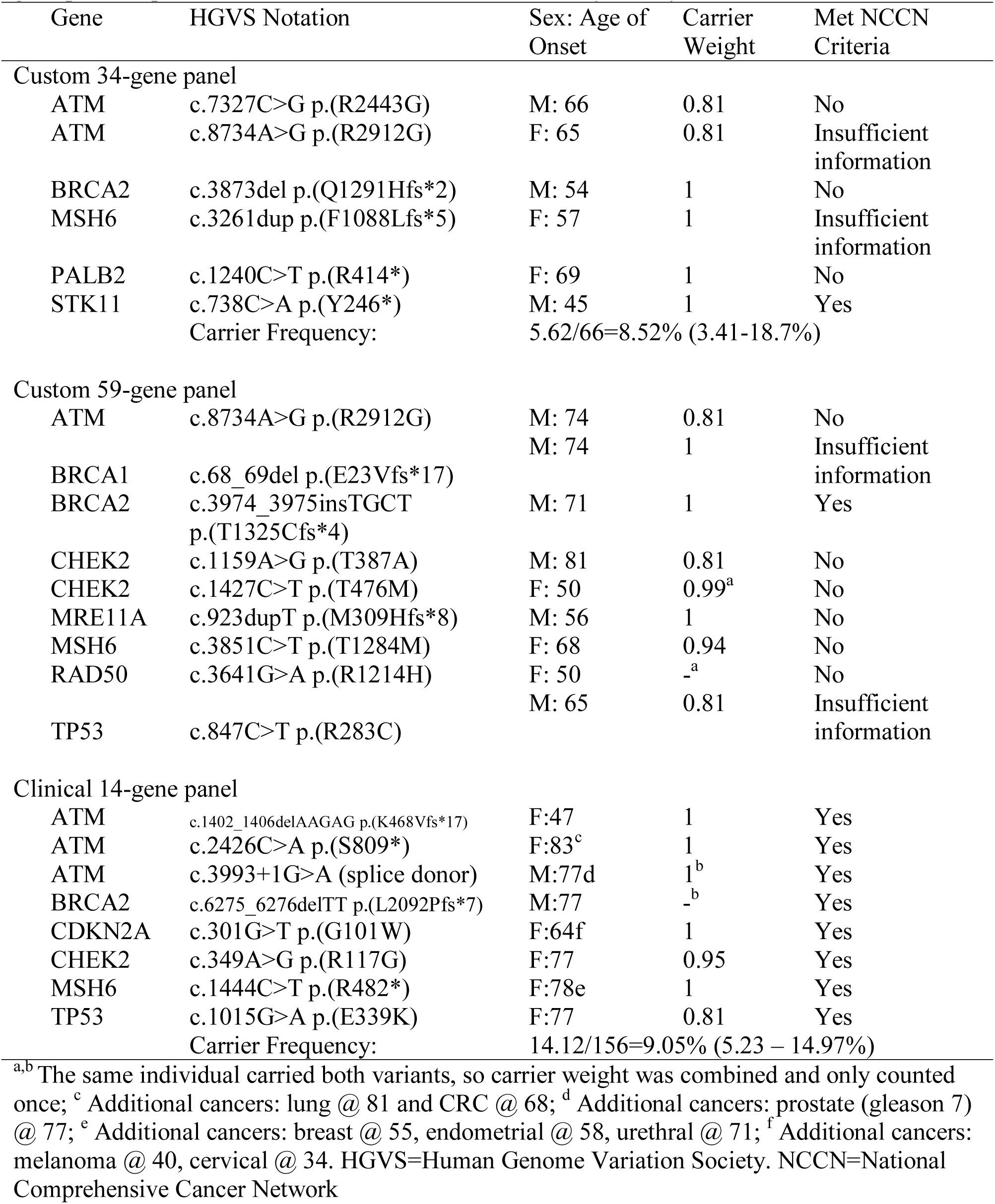
Pathogenic variants and variants of uncertain significance (VUS) with elevated probabilities of pathogenicity identified from a custom 34-or 59-gene panel and a clinical 14-gene panel in pancreatic cancer cases, unselected for family history.

Sequences from the Utah cohort with ≥ 100X mean coverage and 154 pancreatic cancer cases from the Cancer Genome Atlas (TCGA)[26] were analyzed using the USeq (useq.sourceforge.net) in-house pipeline, according to the Genome Analysis Toolkit (GATK v.3.3-0) best practices recommendations.[27] Variants with a mapping quality score less than 20 were excluded. ANNOVAR was used for variant functional annotation followed by conversion to Human Genome Variation Society (HGVS) nomenclature using Mutalyzer.[28, 29]

### Sequence variant evaluation

Truncating variants not present in the final exon of a gene were considered pathogenic. The following filters were used to exclude variants from further analysis: minor allele frequency ≥ 0.1% in one or more populations from the Exome Aggregation Consortium (ExAC) database;[30] synonymous/intronic variants with no predicted effect on splicing via MaxEntScan;[31] variants reported as probable-non-pathogenic/non-pathogenic by more than one source with no conflicting reports in ClinVar (www.ncbi.nlm.nih.gov/clinvar).

Variants of uncertain significance (VUS) were included if their estimated prior probabilities of pathogenicity were >0.8 based on calibrated *in silico* predictions from publicly available databases for the mismatch repair (MMR) genes (hci-lovd.hci.utah.edu), or *BRCA1*/2 (http://priors.hci.utah.edu/PRIORS/). VUS of this type were weighted according to their sequence analysis-based prior probability of pathogenicity (Prior_P) score. The remaining rare VUS were included if at least three of the four missense analysis programs Align-GVGD, MAPP, Polyphen-2, and CADD predicted a severe score.[32–36] This filter corresponds with an OR=3.27 when comparing early-onset breast cancer cases with matched controls.[35] Based on the likelihood ratios identified for *BRCA1*/2,[37] this grouping was assigned a weight of 0.81; from here forward, we refer to these as high probability of pathogenicity VUS (HiP-VUS). Canonical splice acceptor/donor variants predicted to impact splicing were given the weight of 0.97 if the effect of the variant had not been demonstrated experimentally.[38] Pathogenic variants and HiP-VUS detected by the 34-/59-gene panels were confirmed via Sanger sequencing. VUS reported by the Mayo Clinic[39] plus the non-TCGA ExAC (excluding the Finnish and undescribed populations), were graded with the same weights and severity to generate a bioinformatically equivalent set of HiP-VUS. An overview of the datasets and methods used for evaluation is shown (Figure 1).

**Figure 1.**
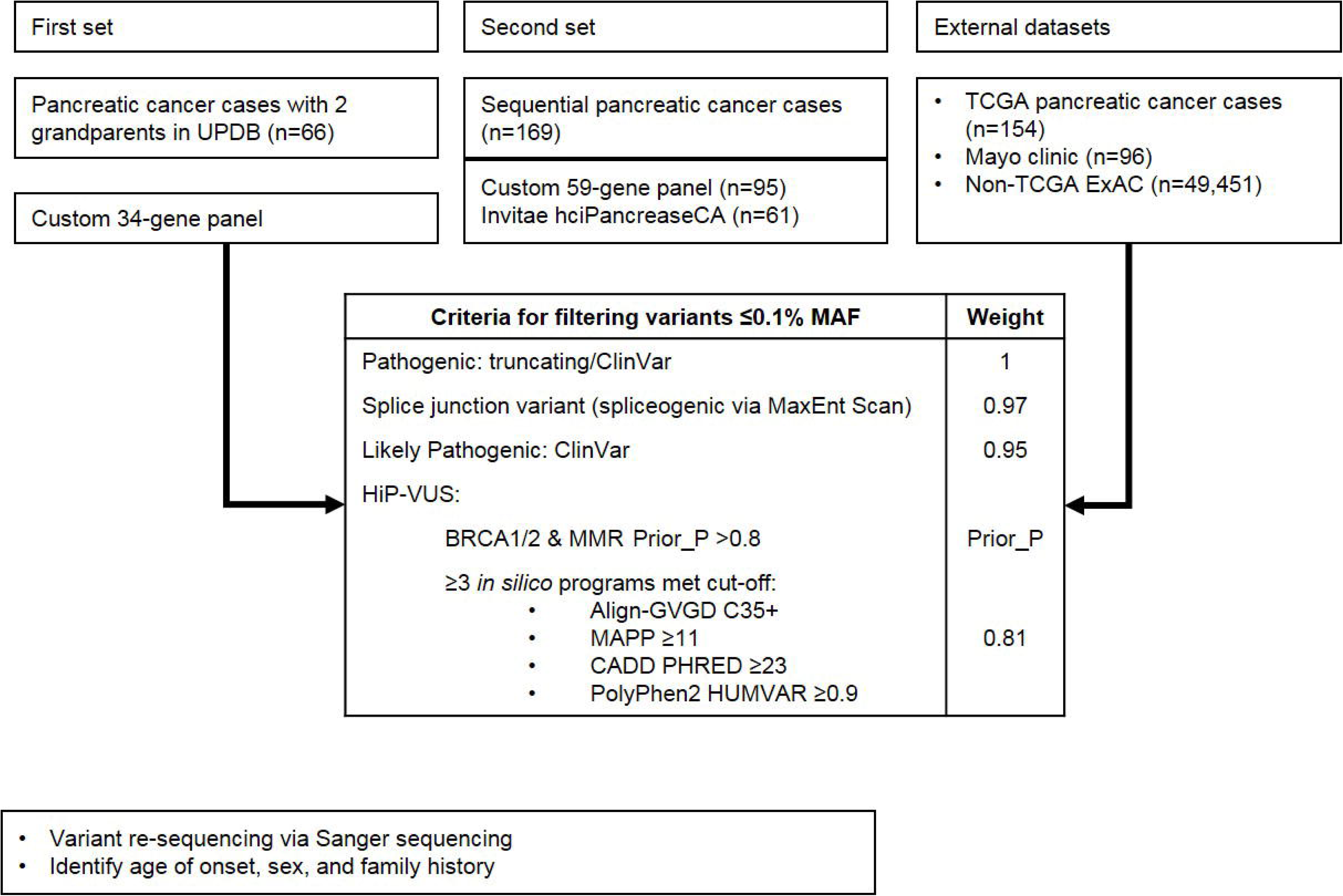
Flowchart of methods. Includes the weighting assigned to filtered sequence variants. In the second set of unselected pancreatic cancer cases, one case failed testing with the custom 59-gene panel and 12 cases declined testing with the Invitae hciPancreasCA panel. MAF = minor allele frequency; HiP-VUS = high probability of pathogenicity variant of uncertain significance; MMR – mismatch repair; Prior_P = sequence analysis-based prior probability of pathogenicity.

### Statistical analysis

STATA V.13.1 (StataCorp, College Station, Texas, USA) was used to conduct meta-analyses, and calculate carrier percentages and 95% confidence intervals. The meta-analyses to compare the carrier frequencies between different pancreatic cancer cohorts were conducted using Metaprop under a random effects model, and Freeman-Tukey transformation to stabilize the variances over the studies.[40] The weighted proportions of variant carriers in the unselected pancreatic cancer cases were compared to the corresponding proportions in the non-TCGA ExAC population to estimate Standardized Incidence Ratios (SIR).[41] Tests of significance and confidence intervals were estimated based on a Poisson distribution.[42] For the meta-analysis and SIR calculation, the genes were split into subgroups of high- and moderate-risk cancer susceptibility genes. High- and moderate-risk were defined as genes with a cumulative risk at age 80 >32% or between 19-32%, respectively, for the cancer with which they are most closely associated.[2] The R package ggplot2 was used to plot the meta-analyses and SIRs.[43]

## RESULTS

### Pancreatic cancer, unselected for family history

In an initial set of 66 pancreatic cancer cases unselected for family history of cancer, 4 pathogenic variants were identified in *BRCA2, MSH6, PALB2*, and *STK11*. After filtering VUS to those with elevated probabilities of pathogenicity, 2 HiP-VUS in *ATM* remained (Table 1). After weighting, 8.5% of these pancreatic cancer cases carried a variant with potential medical management impact for relatives.

To replicate this observation, 156 independent pancreatic cancer cases underwent hereditary cancer predisposition multigene panel testing (n=96 tested with the 59-gene research panel of which one sample failed, and n=61 tested with the 14-gene clinical panel after 12 declined testing). Ten pathogenic variants and seven HiP-VUS were identified (Table 1). These were all identified in genes included in the 34-gene panel. In addition to *in silico* predictions, *CHEK2* p.(T476M) was found to be damaging in a functional assay for *CHEK2* variants and was weighted more strongly towards being pathogenic.[44] After weighting carriers, we found 9.1% of the replication series of pancreatic cancer cases, unselected for family history, carried a variant with potential medical impact.

### Post-variant evaluation of genetic testing eligibility

In order to determine if the pancreatic cancer cases would have qualified for genetic testing, we compared the pancreatic cancer cases that carried pathogenic variants or HiP-VUS with their self-reported family history to published NCCN guidelines.[45] For our initial 66 pancreatic cancer cases, we also were able to access their family history of cancer from UPDB (Supplemental Figure 1). Outside of the STK11 carrier’s family (who had a clinical diagnosis of Peutz-Jegher syndrome), in which the mother and sister had previously been diagnosed, none of the biological relatives had undergone cascade genetic testing.

### Meta-analysis of carrier proportions across studies

The two HCH sets of pancreatic cancer cases were combined with a published study of unselected pancreatic cancer cases from the Mayo Clinic (n=96),[39] plus the pancreatic cancer cases from TCGA (n=154), in a meta-analysis (Figure 2; Supplemental Table 3; Supplemental Table 4). Among unselected pancreatic cancer cases, 4.1% (p=4.9x10^-13^) carried a clearly pathogenic variant in a high-risk cancer susceptibility gene which includes pancreatic cancer in its tumor spectrum.[11] Weighed inclusion of HiP-VUS increased the proportion to 6.0% (p<1.0x10^-16^). For the moderate-risk homologous recombination repair (HRR) breast cancer genes *ATM, BARD1, CHEK2*, and *NBN*,[2, 46] 3.8% (p=8.0x10^-04^) of unselected pancreatic cancer cases carried a clearly pathogenic variant, and weighted inclusion of HiP-VUS increased the proportion to 5.3% (p=2.8x10^-09^). Including all the high-risk genes and the moderate-risk HRR breast cancer genes, 8.1% (p=2.0x10^-14^) of unselected pancreatic cancer cases carry a clearly pathogenic variant, and 10.7% (p<1.0x10^-16^) carry either a clearly pathogenic variant or weighted HiP-VUS with elevated probability of pathogenicity that could enable the at-risk relatives to qualify for preventive HBOC or CRC measures.

**Figure 2.**
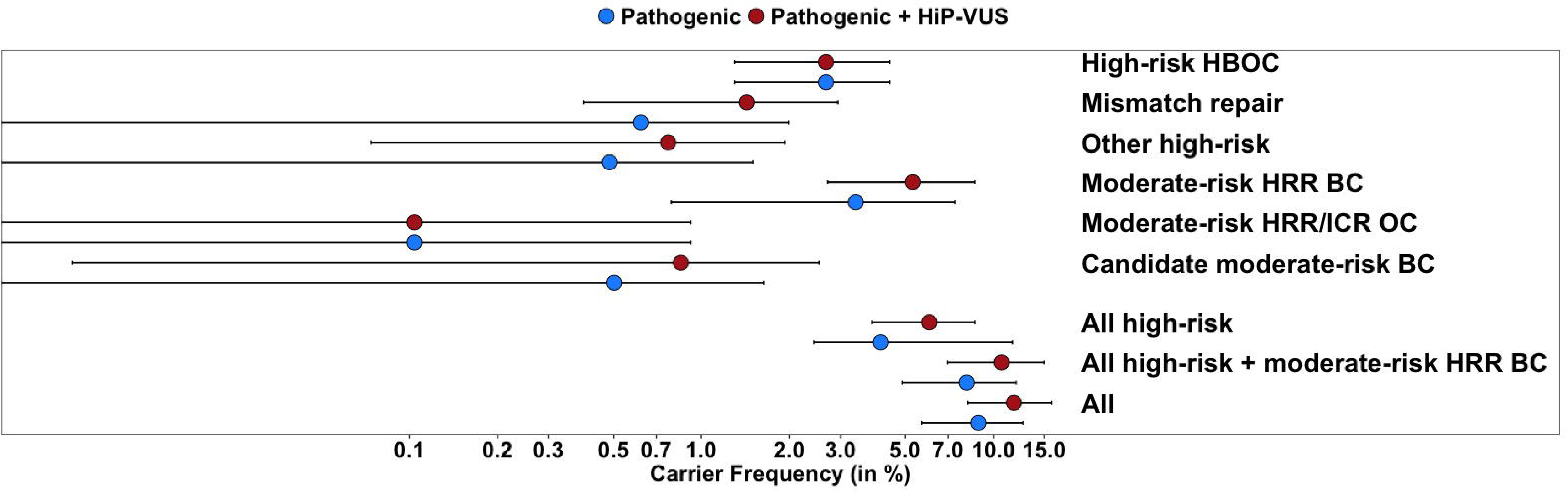
The proportion of carriers of pathogenic variants and high probability of pathogenicity variants of uncertain significance (HiP-VUS) in unselected pancreatic cancer. Results based on a meta-analysis of the unselected pancreatic cancer cases from the Huntsman Cancer Hospital (HCH), the Mayo Clinic, and the pancreatic cancer cases from The Cancer Genome Atlas (TCGA). Carrier frequency point estimates and 95% confidence intervals for groups of genes are presented on a log-scale. A list of genes contained within each analysis group is provided in Supplemental Table 1. The breakdown of results by study is described in Supplemental Table 3. HBOC = Hereditary Breast and Ovarian Cancer; HRR = Homologous Recombination and Repair; ICR = Interstrand Crosslink Repair; OC = Ovarian Cancer; BC = Breast Cancer.

Further, the gene burdens observed in the Utah, Mayo, and TCGA were compared to the non-TCGA ExAC (n=49,451, excluding the Finnish and other subpopulations) as a population sample to determine SIR for subgroups of genes (Figure 3, Supplemental Table 5). As a group, the high-risk susceptibility genes had a SIR = 3.1 (p=2.1x10^-05^). The moderate-risk HRR genes had a slightly lower SIR = 2.4 (p=9.2x10^-05^).

**Figure 3.**
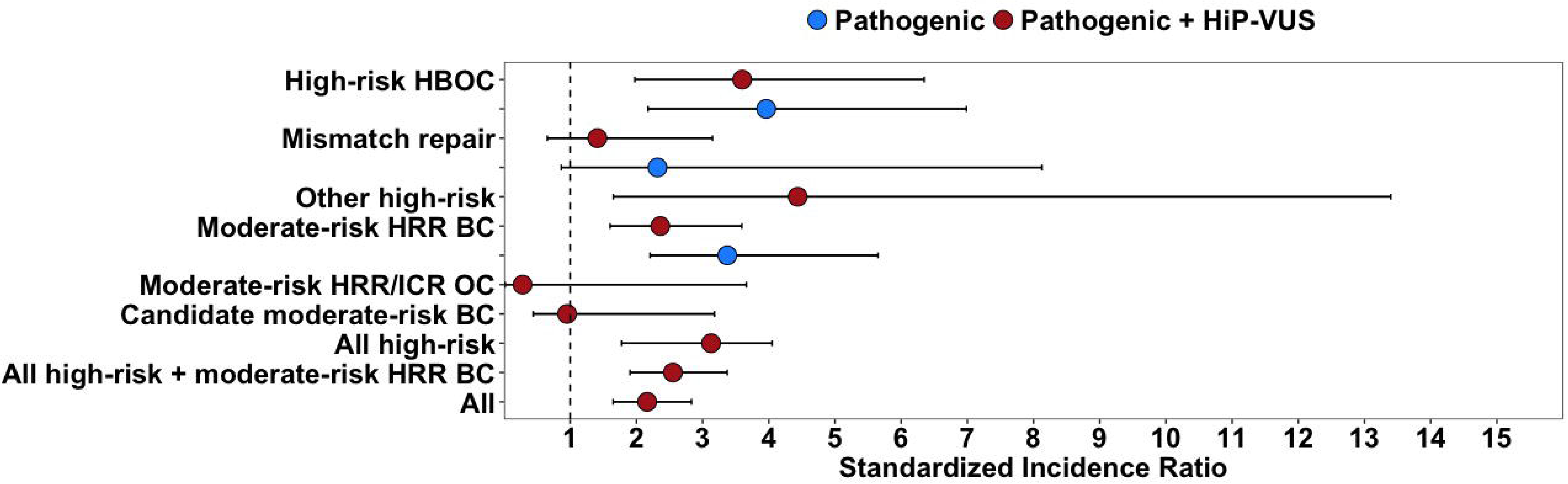
Standardized incidence ratios for cancer susceptibility gene groups in unselected pancreatic cancer cases. The carrier frequencies from the meta-analysis of the cases and the Exome Aggregation Consortium excluding the Cancer Genome Atlas (non-TCGA ExAC) are detailed in Supplemental Table 5. A list of genes contained within each analysis group is provided in Supplemental Table 1. Error bars represent 95% confidence intervals. HBOC =Hereditary Breast and Ovarian Cancer; HRR = Homologous Recombination and Repair; ICR = Interstrand Crosslink Repair; OC = Ovarian Cancer; BC = Breast Cancer.

## Discussion

Through systematic panel testing of pancreatic cancer cases unselected for family history, we estimate that 2.7% (95% CI: 1.3-4.4) carry a pathogenic allele of a high-risk HBOC gene (*BRCA1, BRCA2*, or *PALB2*) and 1.4% (95% CI: 0.4-2.9) of probands carry a likely pathogenic allele of a Lynch Syndrome (LS)-associated MMR gene (*MLH1, MSH2, PMS2*, or *MSH6*). Adding other high-risk genes such as *TP53, CDKN2A*, and *STK11* results in 6.0% (95% CI: 3.9-8.6) of pancreatic cancer cases with a sequence variant that would alter medical management of healthy at-risk relatives: i.e. MRI in addition to mammography or early colonoscopy (Figure 2).[16, 47–50] Additionally, 5.3% (95% CI: 2.7-8.6) carry a likely pathogenic variant of a moderate-risk breast cancer susceptibility gene (i.e., *ATM, BARD1, CHEK2*, or *NBN*) bringing the total proportion of estimated carriers to 10.7% (95% CI: 7.0-15.0). Here we note that *ATM* and *CHEK2* have recently been added to NCCN’s list of genes with associated medical action for breast cancer.[16] Focusing on individual genes, the top four genes with potential medical impact for at-risk relatives, based on weighted counts, were *ATM* (identified in 4 cases), *BRCA2* (3 cases), *CHEK2* (3 cases), and *MSH6* (3cases).

Do findings presented here justify universal screening of all pancreatic cancer patients using a panel test? The precedent for a universal approach comes from what has been accepted and learned from universal testing of all CRCs for LS. Universal LS testing, with immunohistochemical (IHC) or microsatellite instability (MSI)-based pre-screen of tumors followed by germline testing for indicated individuals, is recommended for newly diagnosed CRC cases.[47, 51] This strategy may soon be overtaken by germline DNA panel testing for LS due to 1) rapid decline of panel testing cost, 2) superiority of specificity and sensitivity, and 3) evidence that pre-screening delays testing, which results in a subsequent ~50% loss in follow-up by patients.[52–59] Indeed, a health economics analysis recently published by Erten et al.[59] concluded that universal testing of CRC patients for LS based on sequencing alone will become more cost-effective than the two-step test when the cost of MMR gene sequencing drops to or below $609 USD, echoing a similar finding by Gould-Suarez et al.[59, 60] Based on our results, universal testing of pancreatic cancer patients using a panel test would identify pathogenic variants or MMR HiP-VUS in 1.4% of pancreatic cancer; this is quite comparable to currently accepted two-step universal testing of CRC patients, which identifies pathogenic MMR variants in about 1.2% of patients.[59]

In a recent study, 11.8% of unselected patients with metastatic prostate cancer were found to carry pathogenic variants in DNA-repair genes.[61] Pritchard et al. suggest that this proportion of metastatic prostate cancer cases is high enough to utilize metastatic prostate cancer as a sentinel for cancer predisposition testing. The 11.8% proportion observed in metastatic prostate cancer is similar to the 10.7% observed in this meta-analysis of pancreatic cancer cases. Since the proportion of carriers are similar for pancreatic and metastatic prostate cancer, both of these areas will need to be re-evaluated for guideline updates. For these patients, universal panel testing offers critical time and convenience advantages over cascade testing strategies, resulting in a decreased loss to follow-up or mortality and correspondingly increased benefit to at-risk relatives.

Results obtained here are also relevant to treatment of pancreatic cancer cases with germline pathogenic variants or HiP-VUS in several of the genes included in panel tests. In cells with biallelic pathogenic variants in BRCA1, BRCA2, or PALB2, DNA double strand break repair by homologous recombination (HRR) is badly compromised. Exposure of cells with an HRR deficit to poly (ADP-ribose) polymerase inhibitors (PARPi) creates a synthetic lethal interaction with a wider therapeutic window than many other anti-cancer treatments.[62, 63] As these three genes are most notably associated with breast and ovarian cancer, clinical PARPi studies began with those diseases. Nonetheless, individual patient reports and phase I trial results provide at least anecdotal evidence for beneficial responses in pancreatic cancer patients.[64, 65]

Part of the rationale behind immunotherapy is that mutational processes taking placing during tumor initiation and progression create neoantigens that the patient immune system could respond against. This is true in tumors with compromised HRR, but even more so in tumors with compromised DNA mismatch repair (MMR) which is often caused by biallelic pathogenic variants in MLH1, MSH2, PMS2, or MSH6.[66, 67] A tumor progressed from initiation to clinical relevance implies that it circumvented immune surveillance, immune checkpoint blockade therapies, such as inhibition of PD-1 or PD-L1, designed to derepress immune responses towards a tumor have proven effective in a variety of MMR-deficient solid tumors.[68] Therefore, immunotherapy could be a promising in the ~1.5% of pancreatic tumors arising in patients with LS.

## Acknowledgements

We acknowledge and appreciate the families for their continued support and participation in our studies. This research was supported by United States National Institutes of Health (NIH) National Cancer Institute (NCI) grant R01CA164138; NCI grant P30CA042014 (support of Genetic Counseling Shared Resource), by the Utah Genome Project; by the Canadian Institutes of Health Research (CIHR) for the CIHR Team in Familial Risks of Breast Cancer program; by the Government of Canada through Genome Canada and the Canadian Institutes of Health Research, and the Ministère de l’enseignement supérieur, de la recherche, de la science, et de la technologie du Québec through Génome Québec; by NIH NCI Cancer Center Support Grant P30CA042014; and by the Huntsman Cancer Foundation. The results published here are in part based upon data generated by The Cancer Genome Atlas managed by the NCI and NHGRI. Information about TCGA can be found at http://cancergenome.nih.gov. BAT is a National Health and Medical Research Council CJ Martin Early Career Fellow.

## Notes

**Conflict of Interest Statement**: none to declare

